# Flux balance analysis predicts NADP phosphatase and NADH kinase are critical to balancing redox during xylose fermentation in *Scheffersomyces stipitis*

**DOI:** 10.1101/390401

**Authors:** Kevin Correia, Anna Khusnutdinova, Peter Yan Li, Jeong Chan Joo, Greg Brown, Alexander F. Yakunin, Radhakrishnan Mahadevan

**Affiliations:** Department of Chemical Engineering and Applied Chemistry, University of Toronto, Canada, ON; Institute of Biomaterials and Biomedical Engineering, University of Toronto, Ontario, Canada

## Abstract

Xylose is the second most abundant sugar in lignocellulose and can be used as a feedstock for next-generation biofuels by industry. *Saccharomyces cerevisiae*, one of the main workhorses in biotechnology, is unable to metabolize xylose natively but has been engineered to ferment xylose to ethanol with the xylose reductase (XR) and xylitol dehydrogenase (XDH) genes from *Scheffersoymces stipitis*. In the scientific literature, the yield and volumetric productivity of xylose fermentation to ethanol in engineered *S. cerevisiae* still lags *S. stipitis*, despite expressing of the same XR-XDH genes. These contrasting phenotypes can be due to differences in *S. cerevisiae’s* redox metabolism that hinders xylose fermentation, differences in *S. stipitis’* redox metabolism that promotes xylose fermentation, or both. To help elucidate how *S. stipitis* ferments xylose, we used flux balance analysis to test various redox balancing mechanisms, reviewed published omics datasets, and studied the phylogeny of key genes in xylose fermentation. *In vivo* and *in silico* xylose fermentation cannot be reconciled without NADP phosphatase (NADPase) and NADH kinase. We identified eight candidate genes for NADPase. *PHO3.2* was the sole candidate showing evidence of expression during xylose fermentation. Pho3.2p and Pho3p, a recent paralog, were purified and characterized for their substrate preferences. Only Pho3.2p was found to have NADPase activity. Both NADPase and NAD(P)H-dependent XR emerged from recent duplications in a common ancestor of *Scheffersoymces* and *Spathaspora* to enable efficient xylose fermentation to ethanol. This study demonstrates the advantages of using metabolic simulations, omics data, bioinformatics, and enzymology to reverse engineer metabolism.

## 1 INTRODUCTION

Xylose is the second most abundant sugar in lignocellulose and can be used as a carbon source for biofuels and biochemicals by industry (Jeffries and Jin, 2004). Known catabolic pathways include xylose isomerase (XI) (Schellenberg et al., 1984), the xylose reductase (XR)-xylitol dehydrogenase (XDH) pathway (Horitsu et al., 1968), the Weimberg pathway (Weimberg, 1961), and the Dahms pathway (Dahms, 1974). *Saccharomyces cerevisiae*, one of the main workhorses of industrial biotechnology, has evolved to ferment glucose to ethanol rapidly but cannot natively grow on xylose or ferment it to ethanol (Jeffries and Jin, 2004), even though it has XR and XDH genes. Screening of various yeasts has found that ethanol fermentation with xylose in yeasts is rare (Toivola et al., 1984) and requires NADH-linked XR to alleviate the cofactor imbalance in the XR-XDH pathway during oxygen limitation (Schneider et al., 1981; Slininger et al., 1982; Bruinenberg et al., 1984).

Early research on xylose fermentation in yeasts focused on process optimization of native xylose fermenters (Slininger et al., 1985, 1990; du Preez, 1994), but advancements in genetic engineering enabled the expression of bacterial XI (Sarthy et al., 1987; Amore et al., 1989) and the XR-XDH pathway in *S. cerevisiae* (Kötter and Ciriacy, 1993). *S. cerevisiae* engineered with the XR-XDH pathway from *S. stipitis*, encoded by *XYL1* and *XYL2*, have typically outperformed strains expressing bacterial XI. More recently, evolved strains of *S. cerevisiae* expressing fungal XI has led to faster growth rates and fewer byproducts (Zhou et al., 2012; Verhoeven et al., 2017). Kwak and Jin (2017) provide a review of engineering xylose fermentation in *S. cerevisiae*. After more than 30 years of engineering xylose fermentation in *S. cerevisiae*, the yield and volumetric productivity of engineered *S. cerevisiae* with XI or XR-XDH have lagged native xylose fermenters like *S. stipitis* and *S. passalidarum* in the scientific literature (van Vleet and Jeffries, 2009; Kim et al., 2013a). The inability of *S. cerevisiae*, engineered the XR-XDH pathway from *S. stipitis*, to anaerobically ferment xylose to ethanol at high yields is especially puzzling because the same genes appear to enable xylose fermentation in wild-type *S. stipitis* (Wahlbom et al., 2003). These contrasting phenotypes can be due to differences in the transcriptome, proteome, and metabolome of *S. cerevisiae* that hinder xylose fermentation, differences in the transcriptome, proteome, and metabolome of *S. stipitis* that promote xylose fermentation, or a combination of both. The yeast research community has largely focused on targets in *S. cerevisiae* (van Vleet et al., 2008; Wei et al., 2013; Kim et al., 2013b), while few studies have probed for targets in *S. stipitis* beyond central metabolism (Jeppsson et al., 1995; Freese et al., 2011; Wohlbach et al., 2011). Although the XI pathway has less technical challenges for industrial fermentation than the XR-XDH pathway in *S. cerevisiae*, the redox balancing of the XR-XDH pathway in *S. stipitis* has not been fully elucidated and may offer insight into new redox balancing strategies.

Flux balance analysis (FBA) is a computational method often used to gain insight into metabolism (Österlund et al., 2013; McCloskey et al., 2013), and is well suited to study redox metabolism in yeasts (Pereira et al., 2016). To date, there are four genome-scale network reconstructions (GENRE’s) of *S. stipitis*, the most widely studied xylose fermenting yeast: iBB814 (Balagurunathan et al., 2012), iSS884 (Caspeta et al., 2012), iTL885 (Liu et al., 2012), and iPL912 (Li, 2012). Our xylose fermentation simulations with the four models led to xylitol accumulation when we forced the *in vitro* XR cofactor selectivity (60% NADPH), removed cytosolic NADP-dependent acetaldehyde dehydrogenase from the metabolic models, which is not encoded in *S. stipitis’* genome (Correia et al., 2017), prevented flux through degradation pathways and considered alternative optima. Failing to reconcile *in silico* and *in vivo* xylose fermentation in *S. stipitis*, we created a consensus GENRE for *S. stipitis* to analyze redox balancing mechanisms during xylose fermentation (Figure 1), and reviewed published omics datasets to guide our understanding of potential flux constraints.

**Figure 1.**
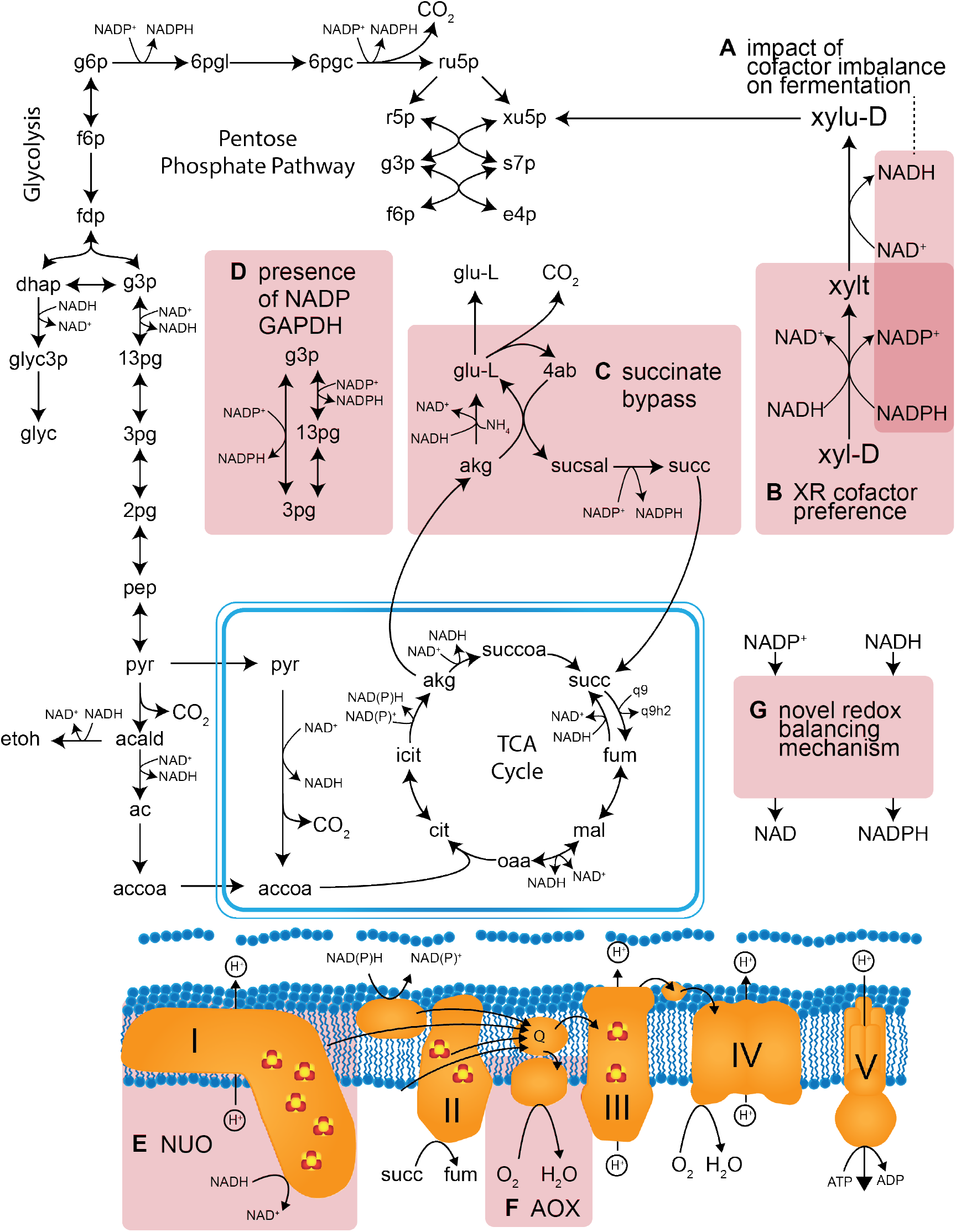
Simplified map of xylose fermentation in *Scheffersomyces stipitis* and potential redox balancing mechanisms. Uncertainites in xylose fermentation are highlighted in red squares: (**A**) the impact of the redox cofactor imbalance on metabolism, (**B**) the *in vivo* XR cofactor preference, (**C**) the use of the succinate bypass to regenerate NADPH, (**D**) the presence of non or phosphorylating glyceraldehyde 3-phosphate dehydrogenase (GAPDH) to regenerate NADPH, (**E**) the impact of bypassing Complex I (NUO) during xylose fermentation, (**F**) the ability of alternative oxidase (AOX) to oxidize NADH during xylose fermentation, and (**G**) the presence of novel redox balancing mechanisms.

The only mechanisms that were able to eliminate *in silico* xylitol accumulation during anaerobic xylose fermentation were phosphorylating NADP-dependent glyceraldehyde 3-phosphate dehydrogenase (GAPDH), non-phosphorylating NADP-dependent GAPDH, or NADP phosphatase (NADPase) and NADH kinase. The expression of phosphorylating NADP-dependent GAPDH, encoded by *K. lactis*’ *GDP1* (Verho et al., 2002), in XR-XDH engineered *S. cerevisiae* decreased its xylitol yield and increased its ethanol yield (Verho et al., 2003); however, there is no strong biochemical or bioinformatic evidence of either form of GAPDH in *S. stipitis*. Expression of cytosolic NADH kinase led to an increase in the xylitol yield of XR-XDH engineered *S. cerevisiae* (Hou et al., 2009), but our simulations indicate that NADPase *and* NADH kinase are both required to completely balance redox cofactors.

The presence of NADPase in *S. cerevisiae* or *S. stipitis* is unknown since it is a eukaryotic orphan enzyme. Eight NADPase candidate genes were identified in iSS885 and iPL912, which had functional annotations from KEGG Orthology (Kanehisa et al., 2011) and PathwayTools (Karp et al., 2009), respectively. The genes encoding the NADPase candidates, the XR-XDH pathway, and enzymes related to NADPH regeneration are outlined in Table 1, along with expression data (Yuan et al., 2011; Huang and Lefsrud, 2012) and orthology information (Correia et al., 2017). *PHO3.2* was the most promising candidate since *S. cerevisiae* does not encode any homologs (Table 1), and its expression in *S. stipitis* was confirmed during xylose fermentation via shotgun proteomics (Huang and Lefsrud, 2012). Therefore, we expressed Pho3.2p and Pho3p, its paralog with 77% identity, in *Komagataella phaffii*, purified them via 6xHis tags, and characterized their substrate preferences. We also studied the phylogenetic origin of our proposed mechanism in the *Scheffersomyces-Spathaspora* clade. We propose NADPase and NADH kinase are critical to xylose fermentation in *S. stipitis* since they can balance redox cofactors in the absence of oxygen (Figure 2). In contrast, *S. cerevisiae* requires oxygen to balance redox, and loses CO_2_ when the oxidative pentose phosphate pathway regenerates NADPH.

**Figure 2.**
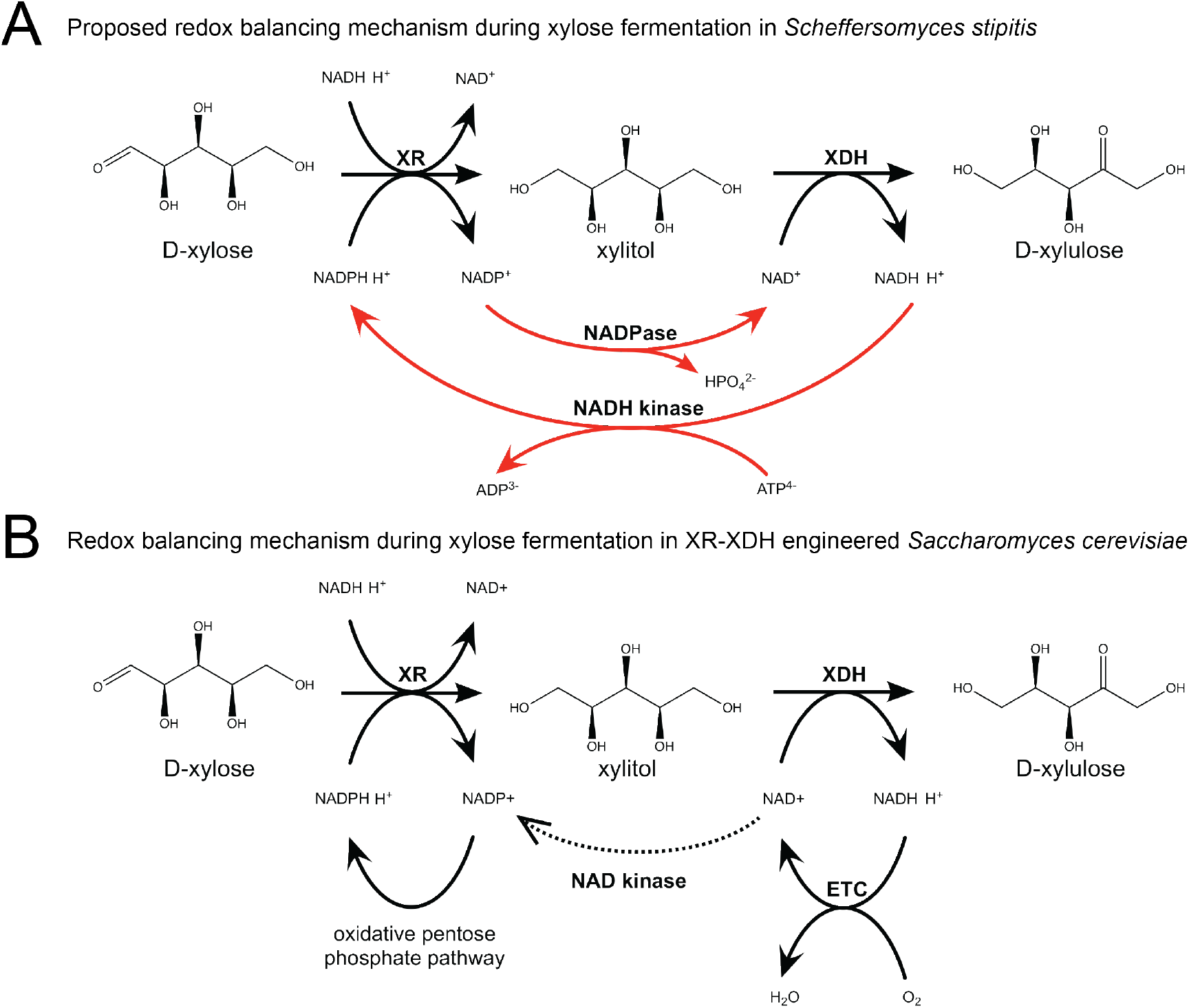
(**A**) Proposed redox balancing during xylose fermentation in *Scheffersomyces stipitis*. NADH kinase regenerates NADPH; NAD(P)H flux drives xylose reductase (XR); NADP phosphatase (NADPase) dephosphylates NADP to NAD; NAD is reduced to NADH by xylitol dehydrogenase (XDH). This redox balancing scheme is consistent with the ^13^C results from (Ligthelm et al., 1988c), independent of oxygen availability, does not have a loss of CO_2_ from the oxidative pentose phosphate pathway, but requires ATP. (**B**) Redox balancing during xylose fermentation in engineered *S. cerevisiae* with the XR-XDH pathway from *S. stipitis*. NAD kinase phosphorylates a fraction of the NAD pool for *de novo* NADP synthesis (dotted line); the oxidative pentose phosphate pathway regenerates NADPH; NAD(P)H drives XR. XDH regenerates NADH; NADH is reoxidized to NAD by the electron transport chain (ETC). Under this redox balancing scheme, there is a loss of CO2 from the oxidative pentose phosphate, oxygen is required to reoxidize NADH, and therefore xylose cannot be anaerobically fermented to ethanol at the maximum theoretical yield.

**Table 1.**
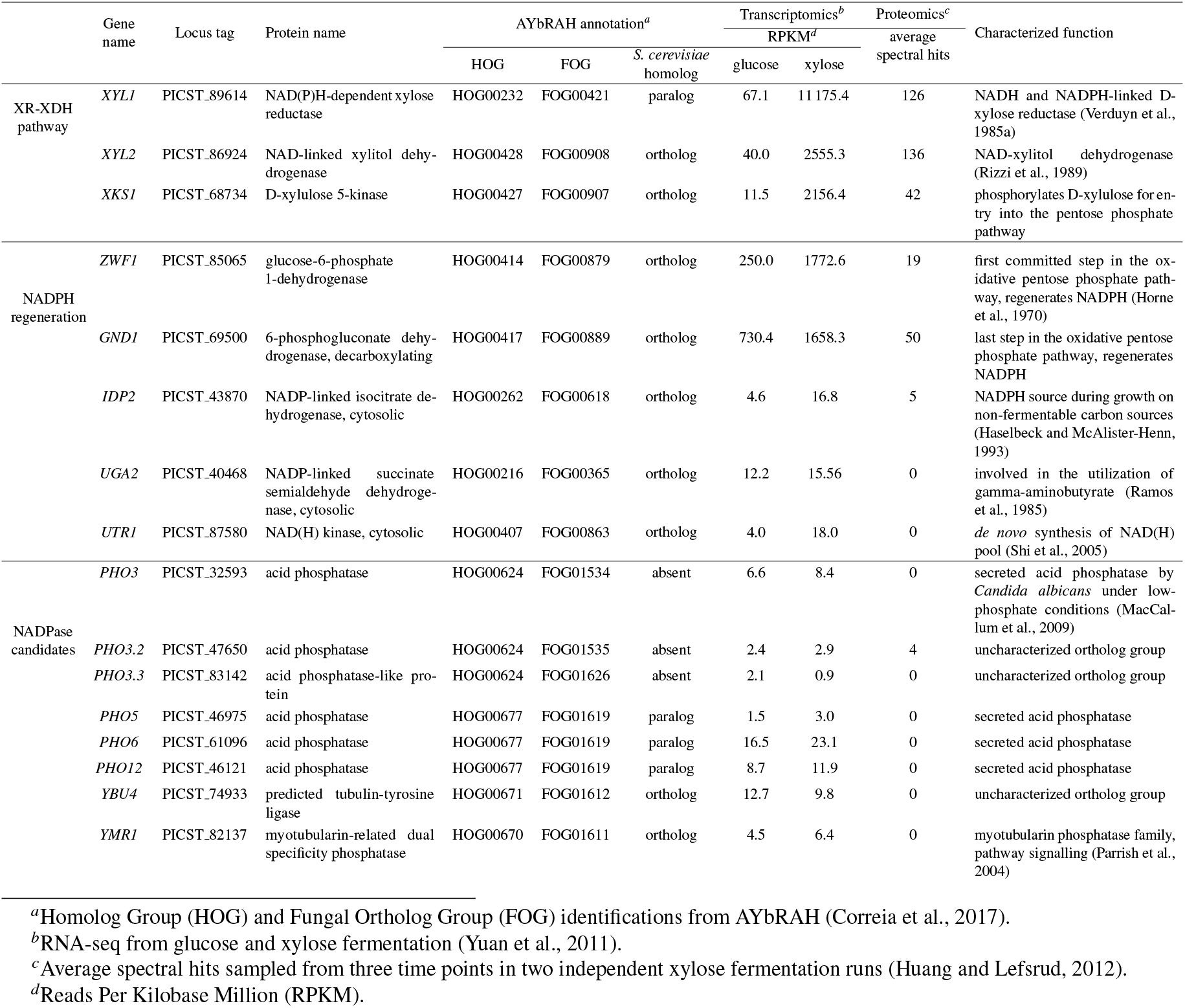
Xylose fermentation related genes in *Scheffersomyces stipitis*, including the xylose reductase(XR)-xylitol dehydrogenase (XDH) pathway, NADPH regeneration, and NADP phosphatase candidates. AYbRAH annotations, transcriptomics, proteomics, and functional characterization are outlined for all the genes.

|  | Gene name | Locus tag | Protein name | AYbRAH annotation<sup>a</sup> |  |  | Transcriptomics<sup>b</sup> |  | Proteomics<sup>c</sup><br>average<br>spectral hits | Characterized function |
| --- | --- | --- | --- | --- | --- | --- | --- | --- | --- | --- |
|  |  |  |  | HOG | FOG | *S. cerevisiae*<br>homolog | glucose | xylose |  |  |
| XR-XDH pathway | *XYL1* | PICST_89614 | NAD(P)H-dependent xylose reductase | HOG00232 | FOG00421 | paralog | 67.1 | 11 175.4 | 126 | NADH and NADPH-linked D-xylose reductase (Verduyn et al., 1985a) |
|  | *XYL2* | PICST_86924 | NAD-linked xylitol dehydrogenase | HOG00428 | FOG00908 | ortholog | 40.0 | 2555.3 | 136 | NAD-xylitol dehydrogenase (Rizzi et al., 1989) |
|  | *XKS1* | PICST_68734 | D-xylulose 5-kinase | HOG00427 | FOG00907 | ortholog | 11.5 | 2156.4 | 42 | phosphorylates D-xylulose for entry into the pentose phosphate pathway |
| NADPH regeneration | *ZWF1* | PICST_85065 | glucose-6-phosphate 1-dehydrogenase | HOG00414 | FOG00879 | ortholog | 250.0 | 1772.6 | 19 | first committed step in the oxidative pentose phosphate pathway, regenerates NADPH (Horne et al., 1970) |
|  | *GND1* | PICST_69500 | 6-phosphogluconate dehydrogenase, decarboxylating | HOG00417 | FOG00889 | ortholog | 730.4 | 1658.3 | 50 | last step in the oxidative pentose phosphate pathway, regenerates NADPH |
|  | *IDP2* | PICST_43870 | NADP-linked isocitrate dehydrogenase, cytosolic | HOG00262 | FOG00618 | ortholog | 4.6 | 16.8 | 5 | NADPH source during growth on non-fermentable carbon sources (Haselbeck and McAlister-Henn, 1993) |
|  | *UGA2* | PICST_40468 | NADP-linked succinate semialdehyde dehydrogenase, cytosolic | HOG00216 | FOG00365 | ortholog | 12.2 | 15.56 | 0 | involved in the utilization of gamma-aminobutyrate (Ramos et al., 1985) |
|  | *UTR1* | PICST_87580 | NAD(H) kinase, cytosolic | HOG00407 | FOG00863 | ortholog | 4.0 | 18.0 | 0 | de novo synthesis of NAD(H) pool (Shi et al., 2005) |
| NADPase candidates | *PHO3* | PICST_32593 | acid phosphatase | HOG00624 | FOG01534 | absent | 6.6 | 8.4 | 0 | secreted acid phosphatase by *Candida albicans* under low-phosphate conditions (MacCallum et al., 2009) |
|  | *PHO3.2* | PICST_47650 | acid phosphatase | HOG00624 | FOG01535 | absent | 2.4 | 2.9 | 4 | uncharacterized ortholog group |
|  | *PHO3.3* | PICST_83142 | acid phosphatase-like protein | HOG00624 | FOG01626 | absent | 2.1 | 0.9 | 0 | uncharacterized ortholog group |
|  | *PHO5* | PICST_46975 | acid phosphatase | HOG00677 | FOG01619 | paralog | 1.5 | 3.0 | 0 | secreted acid phosphatase |
|  | *PHO6* | PICST_61096 | acid phosphatase | HOG00677 | FOG01619 | paralog | 16.5 | 23.1 | 0 | secreted acid phosphatase |
|  | *PHO12* | PICST_46121 | acid phosphatase | HOG00677 | FOG01619 | paralog | 8.7 | 11.9 | 0 | secreted acid phosphatase |
|  | *YBU4* | PICST_74933 | predicted tubulin-tyrosine ligase | HOG00671 | FOG01612 | ortholog | 12.7 | 9.8 | 0 | uncharacterized ortholog group |
|  | *YMR1* | PICST_82137 | myotubularin-related dual specificity phosphatase | HOG00670 | FOG01611 | ortholog | 4.5 | 6.4 | 0 | myotubularin phosphatase family, pathway signalling (Parrish et al., 2004) |

<sup>a</sup>Homolog Group (HOG) and Fungal Ortholog Group (FOG) identifications from AYbRAH (Correia et al., 2017).
<sup>b</sup>RNA-seq from glucose and xylose fermentation (Yuan et al., 2011).
<sup>c</sup>Average spectral hits sampled from three time points in two independent xylose fermentation runs (Huang and Lefsrud, 2012).
<sup>d</sup>Reads Per Kilobase Million (RPKM).

## 2 METHODS

### Genome-scale network reconstruction and analysis

The *S. stipitis* GENRE was obtained from a pan-fungal GENRE that combined the GENRE’s of *S. stipitis* (Balagurunathan et al., 2012; Caspeta et al., 2012; Li, 2012; Liu et al., 2012), *Schizosaccharomyces pombe* (Sohn et al., 2012), *Aspergillus niger* (Andersen et al., 2008), *Yarrowia lipolytica* (Pan and Hua, 2012; Loira et al., 2012), *Komagataella phaffii* (Caspeta et al., 2012), *Kluyveromyces lactis* (Dias et al., 2014), and *S. cerevisiae* (Heavner et al., 2013). Model simulations were carried out using COnstraints Based Reconstruction and Analysis for Python version 0.9.1 (COBRApy) (Ebrahim et al., 2013). The xylose uptake rate was set to 10 mmol · gDCW^−1^ · h^−1^. The growth associated maintenance (GAM) and non-growth associated maintenance (NGAM) were set to 60 mmol · gDCW^−1^ and 0 mmol · gDCW^−1^ · h^−1^, respectively. Flux variability analysis (FVA) was used to evaluate alternative optimum solutions (Mahadevan and Schilling, 2003). The exchange bounds for erythritol, ribitol, arabitol, sorbitol, and glycerol were all set to zero to simplify the solution space for polyols. The cofactor selectivities of NADH and NADPH were varied in a single XR reaction (Balagurunathan et al., 2012). XR solely driven by NADH or NADPH were blocked. The ethanol yield or biomass growth rate were maximized depending on the simulation. We reviewed the data from all published transcriptomics and proteomics studies for *S. stipitis* to guide our understanding of its metabolism (Jeffries et al., 2007; Jeffries and Van Vleet, 2009; Wohlbach et al., 2011; Yuan et al., 2011; Huang and Lefsrud, 2012; Papini et al., 2012; Huang and Lefsrud, 2014).

### Cloning *PHO3/PHO3.2*, and transformation in *K. phaffii*

*PHO3* and *PHO3.2* were amplified from *Scheffersomyces stipitis* CBS 5773 genomic DNA with Phusion polymerase (NEB), without their native signal peptides, as predicted by SignalP (Petersen et al., 2011). The genes were inserted into pPICZα,B using restriction enzyme digestion and ligation. Primer sequences can be found in Table S1. The plasmids were transformed into *Escherichia coli* BL21 using electroporation, selected on low salt LB (Lennox)-Zeocin™ (10μg/mL), and sequence verified (the Centre for Applied Genomics, Toronto). The plasmids were linearly digested by PmeI, and 5 μg of each digestion were transformed into *K. phaffii* KM71H using electroporation (EasySelect *Pichia* Expression Kit, Invitrogen). Cells were recovered in 4mL of YPD for 2 hours and plated on YPD agar with Zeocin™ (100 μg/mL).

### Agar acid phosphatase assay

An acid phosphatase assay was used to screen for *K. phaffii* colonies with the highest expression of Pho3p and Pho3.2p (Dorn, 1965). Colonies were plated on BMMY agar and incubated at 30°C. 100 μL of 100% methanol was dispensed on the lid of each inverted Petri dish after 24 and 48 hours of growth. 20 mg of Fast Garnet GBC sulphate salt (MilliporeSigma) and 2 mg of 1-naphthyl phosphate disodium salt (MilliporeSigma) were dissolved in 4mL of 0.6M acetate buffer (pH 4.8). 4 mL of the solution was flooded into the Petri dish and examined for five minutes (Figure S1).

### General phosphatase assay with para-nitrophenyl phosphate (pNPP)

In addition to the agar acid phosphatase assay, a general phosphatase assay was used to monitor the activity of various *K. phaffii* clones for secreted phosphatase in liquid media (Kuznetsova et al., 2005). 100 μL of the fermentation broth from wild-type *K. phaffii* and mutants expressing Pho3p and Pho3.2p were assayed for pNPP phosphatase after 24 hours of induction. The supernatant enriched with phosphatase from each culture were prepared by treatment with 0.1% of Triton X100 or Tween 20 on a rotator at +4°C for 30 minutes or sonication in 1 mL volume during 10 seconds on ice. The supernatant was separated from the cells by centrifugation at 13 000 rpm and a benchtop Eppendorf 5424 centrifuge. The cells were resuspended in an equal volume of BMMY media. The pNPP phosphatase assay was carried out in 200 μL reactions in sealed 96 well-plates incubated overnight at 30°C. The reaction mixture consisted of the phosphatase-enriched supernatant collected from the fermentation, 4mM pNPP, 0.5mM MnCl_2_, 5mM MgCl_2_, and 100mM HEPES pH 7.5. pNPP phosphatase activity was estimated due to the increasing absorbance of para-nitrophenyl at 410 nm; the absorbance of wild-type *K. phaffii* cultures was subtracted as background.

### Protein expression and purification

The clones with the highest expression of Pho3p and Pho3.2p were grown in BMGY as described in EasySelect *Pichia* Expression Kit (Invitrogen™). 5 mL of 100% methanol was added at 24 and 48 hours after inoculation in 1L of broth in a 4 L baffled flask. We failed to purify any secreted protein in the fermentation broth after 24, 48 and 72 hours after methanol induction; the final pH was adjusted to 7.5, buffered with 50mM HEPES, 0.4M NaCl and 5mM imidazole for Ni-NTA binding. Most of the phosphatase activity was found to be associated with the cells (Figure S2). The cells were collected and stored at −20°C until purification. The cells were sonicated (Qsonica, dual horn probe, 2.5 minutes, 80% of maximal amplitude) to detach the acid phosphatase from the cell surface. The phosphatases relative sizes were estimated to be 55 kDa with glycosylation on 12% PAAG (Figure S3) and were sequence confirmed by mass spectrometry (Supplementary Information). The phosphatases were further purified with a Ni-NTA agarose (Quiagen) by their 6xHis tag according to manufacturer’s protocol. It should be noted that longer sonication time or Y-PER Yeast Protein extraction reagent (Thermo) application increased contamination with an alcohol dehydrogenase (40 kDa band), identified as *K. phaffii* Adh2p (A0A1B2JBQ8) (Figure S3). Size exclusion chromatography (Superdex 10/300 GL) failed to separate the phosphatases from *K. phaffii* Adh2p. The proteins formed a tight dimer, with a relative size of 110kDa (data not shown). We attempted to purify Utr1p from *S. cerevisie* and *S. stipitis* using *E. coli* and *K. phaffii* as expression hosts, but were unable to collect soluble protein.

### Alcohol dehydrogenase enzyme assay

The *K. phaffii* Adh2p contaminant was purified from wild-type *K. phaffii* and assayed for NADH and NADPH oxidation at 340 nm in 96 well-plates at 30 °C, with 0.01-10mM butyraldehyde, 1 mM ZnCl_2_, 50mM HEPES pH −7.5. Both NADH and NADPH reduced butyraldehyde, but no butanol oxidation with NAD or NADP was detected.

### NADPase phosphatase (NADPase) enzyme assay

NADPase activity was assayed in a coupled reaction with formate dehydrogenase (P33160). 2.9 μg of Pho3p and Pho3.2p were incubated at 30°C in 200 μL reactions with NADP, 50 mM sodium formate, 20 μg of strictly NAD-dependent formate dehydrogenase, 50 mM HEPES pH 7.5, 5 mM MgCl_2_ and 0.5 mM MnCl_2_. The reduction of NAD was monitored at 340 nm in 96-well microplates. 10 mM butyraldehyde was used to estimate the concentration of the contaminated Pho3p and Pho3.2p Ni-NTA eluted samples via NADH-dependent butyraldehyde reductase activity.

### Phosphatase screen with natural substrates using the malachite green assay

Pho3p and Pho3.2p were assayed for their substrate preferences with the malachite green assay (Kuznetsova et al., 2015). The assay was performed in 96-well microplates in 160 μL reactions containing of 100 mM HEPES pH 7.5, 5mM MgCl_2_, 0.5mM MnCl_2_, and 3μg the acid phosphatase. 3.125mM was the final substrate concentration of cytidine 2’ monophosphate, inosine triphosphate, NADP, FMN, coenzyme A, glyphosate, adenosine 3’,5’ diphosphate. The remaining substrates were assayed at 6.25mM: AMP, CMP, GMP, IMP, UMP, XMP, 2’AMP, 2’CMP, 3’AMP, 3’CMP, dAMP, dCMP, dGMP, dIMP, dTMP, dUMP, ADP, CDP, GDP, IDP, TDP, UDP, dADP, dCDP, dGDP, ATP, CTP, GTP, ITP, TTP, UTP, dATP, dCTP, dGTP, dITP, dUTP, NADP, FMN, PEP, CoA, phosphocholine, α-glucose 1-phosphate, β-glucose 1-phosphate, ß-glucose 6-phosphate, fructose 1-phosphate, fructose 6-phosphate, ribose 5-phosphate, mannose 1-phosphate, mannose 6-phosphate, galactose 1-phosphate, fructose 1,6-bisphosphate, erythrose 4-phosphate, trehalose 6-phosphate, glucose 1,6-bisphosphate, sucrose 6-phosphate, 2-deoxy-D-glucose 6-phosphate, 2-D-ribose 5-phosphate, glucosamine 6-phosphate, 6-phospho-D-gluconate, L-2-phosphoglycerate, 3-phosphoglycerate, glyceraldehyde 3-phosphate, phytic acid, thiamine monophosphate, thiamine disphosphate, phosphoserine, phosphothreonine, phosphotyrosine, 2-phosphoascorbate, pyridoxal 5-phosphate, polyphosphate, glycerol 1-phosphate, glycerol 2-phosphate, glycerol 3-phosphate, ribulose 1,5-bisphosphate, lactose 1-phosphate, N-acetyl-α-D-glucosamine 1-phosphate, α-D-glucosamine 1-phosphate, N-acetyl-α-D-glucosamine 6-phosphate, dihydroxyacetone phosphate, phosphono-acetate, phosphono-formate, phosphono-methyl-glycine, AMP-ramidate, D-sorbitol 6-phosphate, glyphosate, phosphoryl-ethanolamine, NMN, disphosphate, 2’,5’-ADP, PAP (3’, 5’ - ADP), PAPS, pNPP, 5-methyl dCMP After 30 minutes of incubation at 30°C, free phosphate concentration was estimated by mixing a reaction aliquot with 40 μL of the malachite green solution to a final volume of 200 μL volume. After 1 minute of 1000 rpm orbital shaking, the production of phosphate was measured according to the optical density at 630 nm. The malachite green solution was prepared fresh by mixing 1 mL of 7.5% NH_4_MoO_4_ with 80 μL 11% Tween-20 and malachite stock solution (1 L: 1.1 g malachite green, 150 mL H_2_SO_4_, 750 mL water).

### Syntenic and phylogenetic analysis of XYL1 and *PHO3.2* homologs

AYbRAH was used to annotate proteins in the *XYL1* and *PHO3.2* loci, since genome annotations are lacking for several *Scheffersomyces* and *Spathaspora* species (Correia et al., 2017). The protein sequences of Xyl1p and Pho3.2p from *S. stipitis* were queried against the genomic nucleotide sequences of *Debaryomyces hansenii, Suhomyces tanzawaensis, Scheffersomyces* species, and *Spathaspora* species using TBLASTN; assembly accessions are listed in Table 2. The genomic loci 50 kbp upstream and downstream of the *XYL1* and *PHO3.2* hits were then queried with a protein sequence of the organism’s closest relative for each Fungal Ortholog Group (FOG) in AYbRAH using BLASTP (Correia et al., 2017). The gene coordiantes were manually reviewed to remove spurious open reading frames (see Supplementary Information). Biopython’s GenomeDiagram was used to illustrate the synteny of the *XYL1* and *PHO3.2* loci (scripts available at https://github.com/kcorreia/) (Cock et al., 2009). The *XYL1* and *PHO3.2* homologs were aligned with MAFFT version 7.24. PhyML version 3.2.0 was used to reconstruct the phylogeny of *XYL1* and *PHO3.2* homologs with 1000 bootstrap replicates (Correia et al., 2017). These trees were used to help distinguish between the *XYL1/XYL1.2* and *PHO3/PHO3.2* paralogs.

**Table 2.**
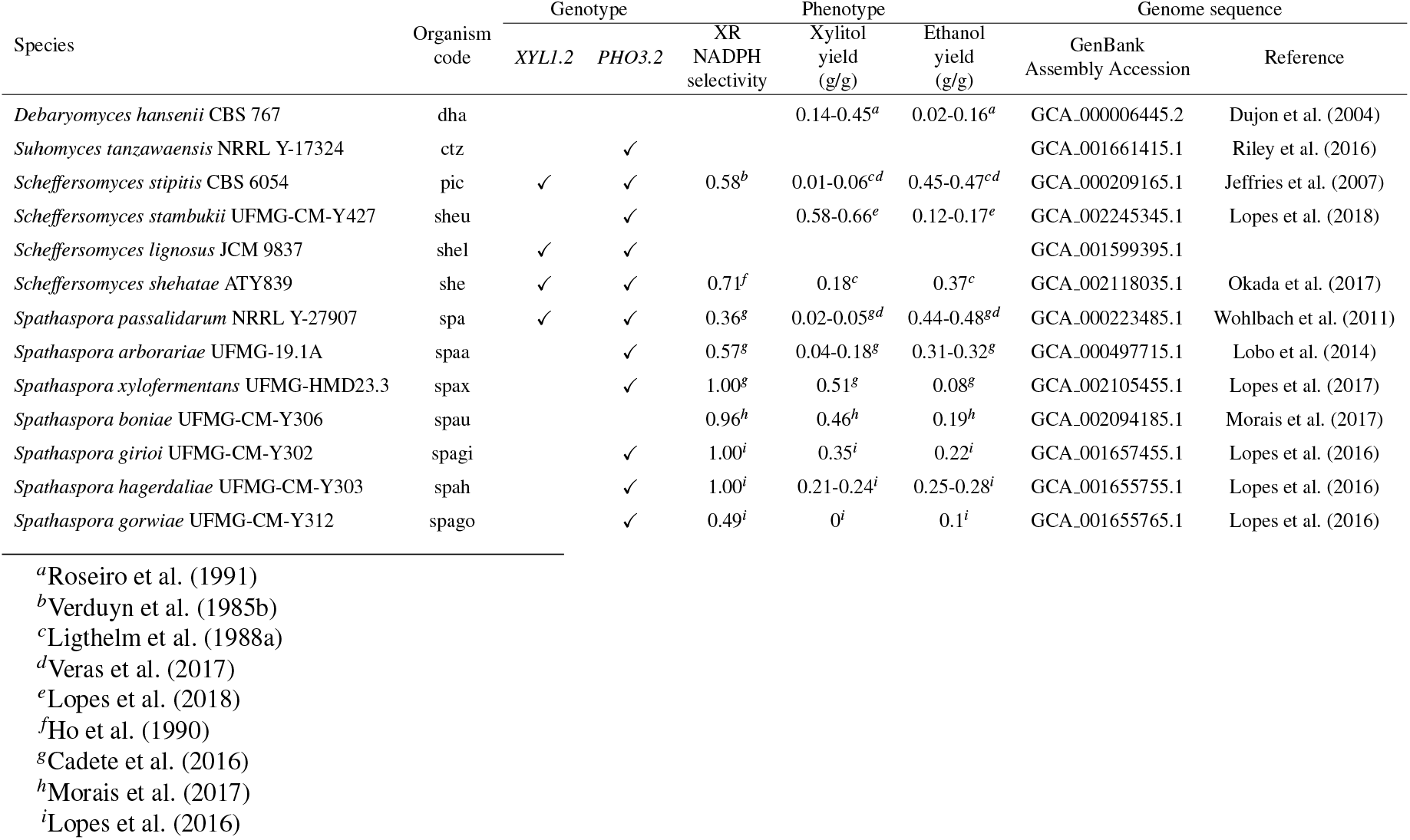
Xylose fermentation genotypes and phenotypes for *Debaryomyces hansenii, Suhomyces tanzawaensis, Spathapora*, and *Scheffersoymces* species. GenBank Assembly Accessions were used to analyze the synteny of *XYL1* and *PHO3.2* loci.

| Species | Organism code | Genotype |  | XR NADPH selectivity | Phenotype |  | Genome sequence |  |
| --- | --- | --- | --- | --- | --- | --- | --- | --- |
|  |  | *XYL1.2* | *PHO3.2* |  | Xylitol yield (g/g) | Ethanol yield (g/g) | GenBank Assembly Accession | Reference |
| *Debaryomyces hansenii* CBS 767 | dha |  |  |  | 0.14-0.45<sup>a</sup> | 0.02-0.16<sup>a</sup> | GCA_000006445.2 | Dujon et al. (2004) |
| *Suhomyces tanzawaensis* NRRL Y-17324 | ctz |  | ✓ |  |  |  | GCA_001661415.1 | Riley et al. (2016) |
| *Scheffersomyces stipitis* CBS 6054 | pic | ✓ | ✓ | 0.58<sup>b</sup> | 0.01-0.06<sup>c,d</sup> | 0.45-0.47<sup>c,d</sup> | GCA_000209165.1 | Jeffries et al. (2007) |
| *Scheffersomyces stambukii* UFMG-CM-Y427 | sheu |  | ✓ |  | 0.58-0.66<sup>e</sup> | 0.12-0.17<sup>e</sup> | GCA_002245345.1 | Lopes et al. (2018) |
| *Scheffersomyces lignosus* JCM 9837 | shel | ✓ | ✓ |  |  |  | GCA_001599395.1 |  |
| *Scheffersomyces shehatae* ATY839 | she | ✓ | ✓ | 0.71<sup>f</sup> | 0.18<sup>c</sup> | 0.37<sup>c</sup> | GCA_002118035.1 | Okada et al. (2017) |
| *Spathaspora passalidarum* NRRL Y-27907 | spa | ✓ | ✓ | 0.36<sup>g</sup> | 0.02-0.05<sup>g,d</sup> | 0.44-0.48<sup>g,d</sup> | GCA_000223485.1 | Wohlbach et al. (2011) |
| *Spathaspora arborariae* UFMG-19.1A | spaa |  | ✓ | 0.57<sup>g</sup> | 0.04-0.18<sup>g</sup> | 0.31-0.32<sup>g</sup> | GCA_000497715.1 | Lobo et al. (2014) |
| *Spathaspora xylofermentans* UFMG-HMD23.3 | spax |  | ✓ | 1.00<sup>g</sup> | 0.51<sup>g</sup> | 0.08<sup>g</sup> | GCA_002105455.1 | Lopes et al. (2017) |
| *Spathaspora boniae* UFMG-CM-Y306 | spau |  |  | 0.96<sup>h</sup> | 0.46<sup>h</sup> | 0.19<sup>h</sup> | GCA_002094185.1 | Morais et al. (2017) |
| *Spathaspora girioi* UFMG-CM-Y302 | spagi |  | ✓ | 1.00<sup>i</sup> | 0.35<sup>i</sup> | 0.22<sup>i</sup> | GCA_001657455.1 | Lopes et al. (2016) |
| *Spathaspora hagerdaliae* UFMG-CM-Y303 | spah |  | ✓ | 1.00<sup>i</sup> | 0.21-0.24<sup>i</sup> | 0.25-0.28<sup>i</sup> | GCA_001655755.1 | Lopes et al. (2016) |
| *Spathaspora gorwiae* UFMG-CM-Y312 | spago |  | ✓ | 0.49<sup>i</sup> | 0<sup>i</sup> | 0.1<sup>i</sup> | GCA_001655765.1 | Lopes et al. (2016) |

<sup>a</sup>Roseiro et al. (1991)
<sup>b</sup>Verduyn et al. (1985b)
<sup>c</sup>Ligthelm et al. (1988a)
<sup>d</sup>Veras et al. (2017)
<sup>e</sup>Lopes et al. (2018)
<sup>f</sup>Ho et al. (1990)
<sup>g</sup>Cadete et al. (2016)
<sup>h</sup>Morais et al. (2017)
<sup>i</sup>Lopes et al. (2016)

## 3 RESULTS & DISCUSSION

### *In vivo* NADPH source in *S. stipitis* during xylose fermentation

*S. stipitis* Xyl1p prefers NADPH to NADH *in vitro*, but its NADPH source and XR selectivity for NADPH are unknown *in vivo*. We first used FVA to analyze the impact of each NADPH source on the maximum ethanol yield from xylose, assuming the *in vitro* NADPH selectivity for XR, in presence and absence of NADPase (Figure 3). The oxidative pentose phosphate pathway is able to ferment xylose to ethanol *in silico*, but its ethanol yields are less than the *in vivo* yields when oxygen uptake rate (OUR) is less than 3 mmol · gDCW^−1^ · h^−1^; the addition of NADPase is unable to resolve the redox imbalance with NADPH regenerated from the oxidative pentose phosphate pathway. Jeffries et al. (2007) proposed the succinate bypass to regenerate NADPH, but it is unable to ferment xylose to ethanol below 10 mmol · gDCW^−1^ · h^−1^ in our simulations. NADP-dependent isocitrate dehydrogenase can also enable ethanol fermentation, but not below 10 mmol · gDCW^−1^ · h^−1^. Only NADPase and NADH kinase, phosphorylating NADP-dependent GAPDH, and non-phosphorylating NADP-dependent GAPDH were able to anaerobically ferment xylose to ethanol at the maximum theoretical yield. There is no strong evidence for NADPH regeneration from NADP-dependent GAPDH in *S. stipitis*, since it does not have any genes orthologous to *K. lactis’* phosphorylating NADP-dependent GAPDH (Correia et al., 2017), and there is no significant upregulation of *UGA2*, the most likely source of nonphosphoryaling GAPDH in *S. stipitis* (Brunner et al., 1998), in its xylose-fermenting transcriptome (Jeffries and Van Vleet, 2009; Yuan et al., 2011). Therefore, these simulations and omics data support NADPase/NADH kinase balancing redox cofactors during xylose fermentation in *S. stipitis* if XR prefers NADPH *in vivo*.

**Figure 3.**
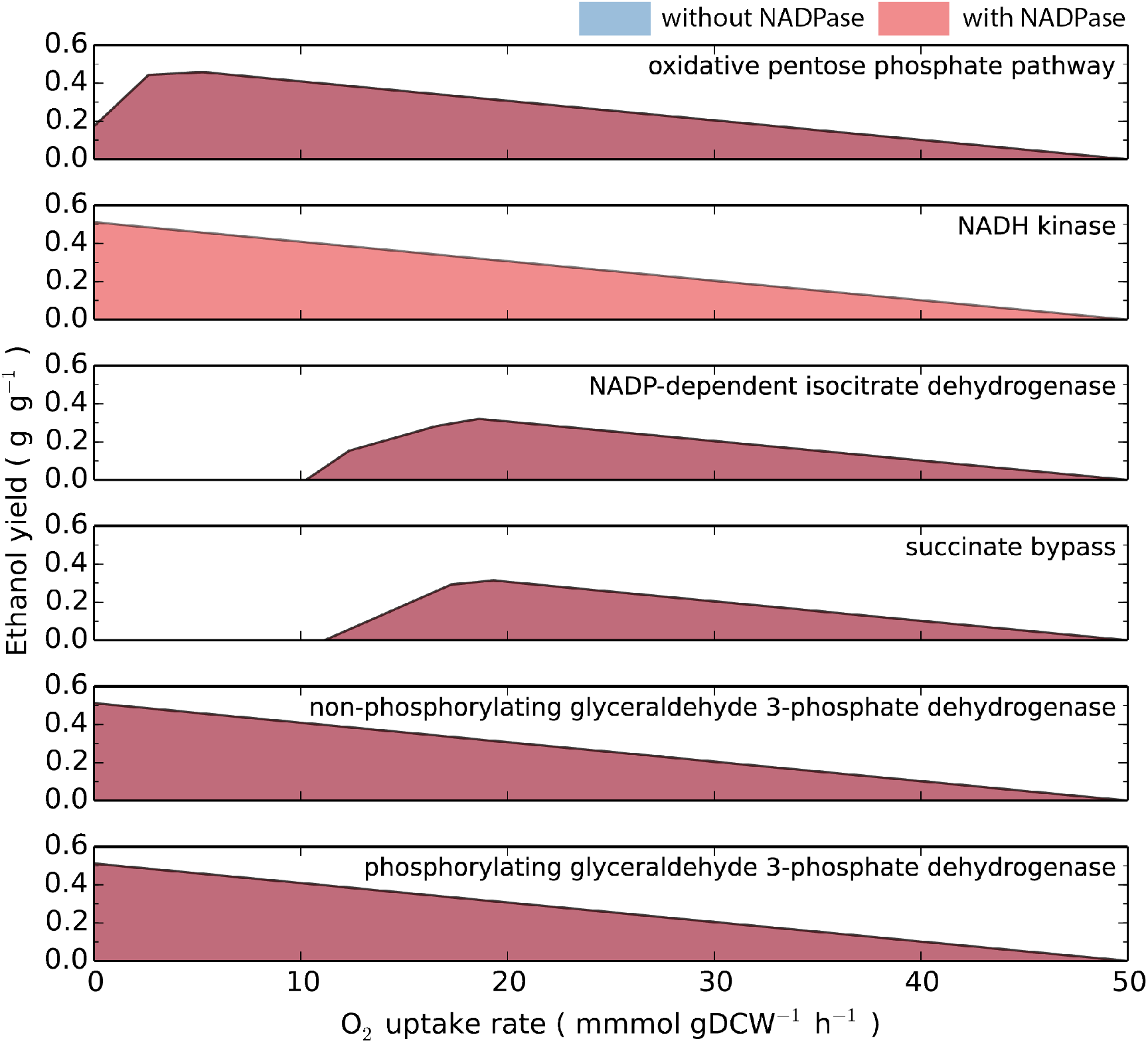
Ethanol yield as a function of NADPH source and oxygen uptake rate (OUR). There is a drop in the ethanol yield when the oxidative pentose phosphate regenerates NADPH at OUR’s close to anaerobic levels. The highest ethanol yields were obtained with NADP phosphatase/NADH kinase, phosphorylating glyceraldehyde 3-phosphate dehydrogenase (GAPDH), and non-phosphorylating GAPDH. NADP-dependent isocitrate dehydrogenase and the succinate bypass were unable to ferment xylose to ethanol below 10 mmol · gDCW^−1^ · h^−1^.

### Impact of XR cofactor preference on xylose fermentation

The *in vivo* XR cofactor preference is not known, so we used FVA to explore the maximum xylitol yield with the ethanol yield as the objective function, with varying OUR’s and NADPH selectivities, in the presence and absence of NADPase/NADH kinase (Figure 4A). Anaerobic xylose fermentation leads to xylitol accumulation when any amount of NADPH drives XR flux *in silico*. The *in silico* xylitol yield falls within the experimental polyol range when the NADPH selectivity is less than 10% NADPH, far less than the XR’s *in vitro* selectivity (60% NADPH). *In silico* anaerobic xylose fermentation to ethanol is infeasible when the XR selectivity for NADPH is greater than 80% in the absence of NADPase and NADH kinase. The *in vitro* cofactor selectivity predicts the xylitol yield to be greater than 0.3 g/g when the OUR is less than 2 mmol · gDCW^−1^ · h^−1^, which is more than the typical polyol yield of 0.1 g/g observed with *S. stipitis* (Ligthelm et al., 1988b; Su et al., 2015). The presence of NADPase/NADH kinase in the model enables the fermentation of xylose to ethanol at all OUR’s and all NADPH selectivities. There is no impact on the ethanol yield when they are present individually. These simulations indicate that either XR can change its cofactor selectivity to NADH during oxygen-limiting conditions, NADPase/NADH kinase plays a role in redox cofactor balancing in *S. stipitis* or additional enzymes are absent in the metabolic model.

**Figure 4.**
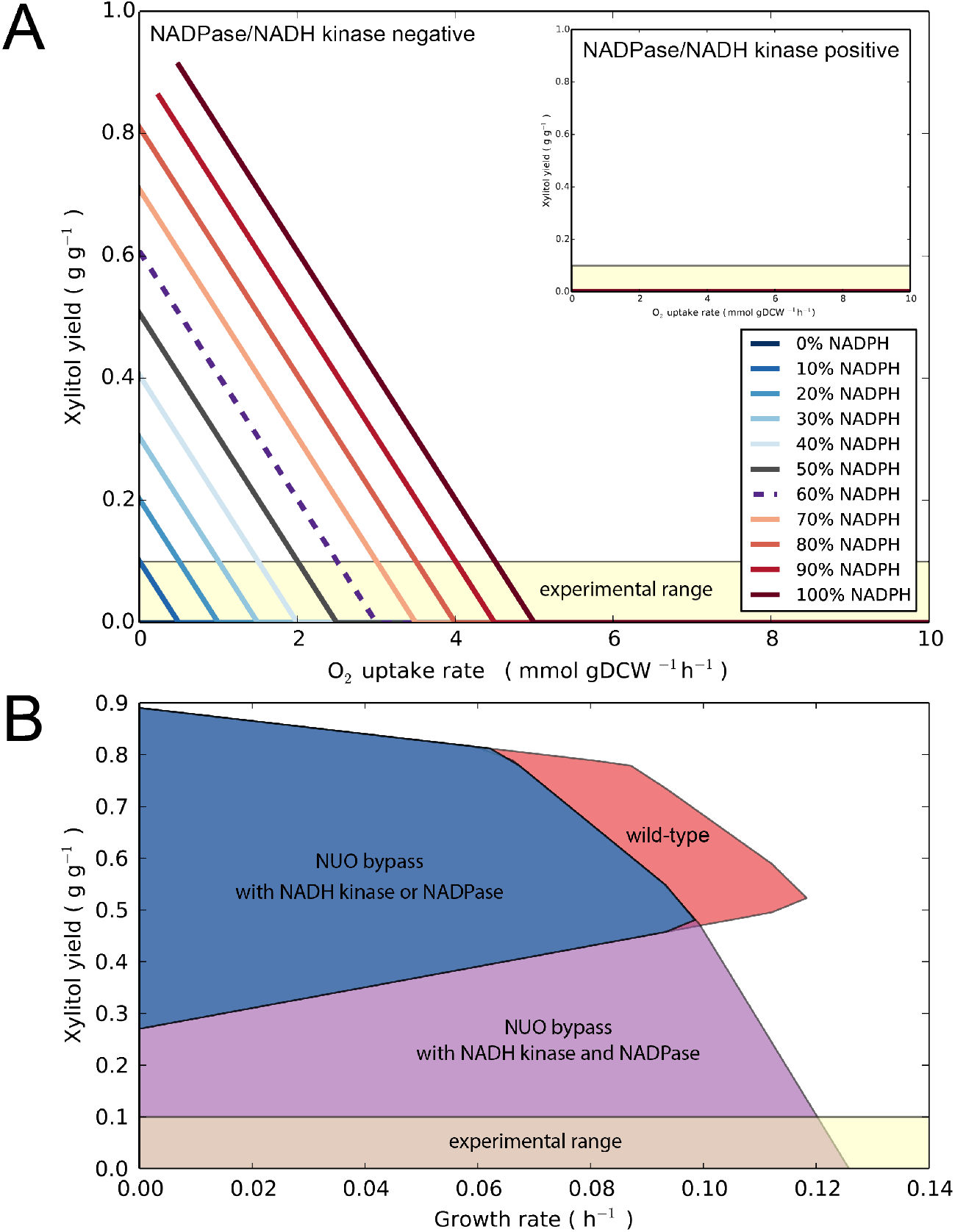
Xylitol yield sensitivity to oxygen uptake rate (OUR) and growth rate. (**A**) Simulations maximized xylose to ethanol with and without NADP phosphatase (NADPase) and NADH kinase. Anaerobic xylose fermentation is only feasible *in silico* when xylose reductase (XR) is driven by more than 80% NADPH. The *in silico* xylitol yield exceeds the 10% polyol yield typically observed *in vivo* when OUR is less than 2 mmol · gDCW^−1^ · h^−1^. The presence of NADPase and NADH kinase in the metabolic model eliminates xylitol yield at all OUR’s and XR cofactor selectivities. (**B**) Xylitol production envelope with and without NADPase and NADH kinase when XR is driven by 60% NADPH. OUR was constrained to 1 mmol · gDCW^−1^ · h^−1^. Simulations without NADPase/NADH kinase lead to xylitol accumulation at the optimal growth rate and at all suboptimal growth rates. Bypassing Complex I (NUO) only reduces the maximum growth rate and has a marginal decrease in the xylitol yield at the optimal growth rate. The presence of NADH kinase or NADPase does not reduce the xylitol yield to the experimental polyol range; however, the addition of NADPase *and* NADH kinase enables the xylitol yield to fall within the experimental polyol range.

Previous attempts to determine the XR cofactor preference are summarized in Table 3. *In vitro* enzyme characterization and crude enzyme assays for XR at varying aeration rates with *S. stipitis* both show higher selectivity for NADPH than NADH. Verduyn et al. (1985b) found that XR preferred NADPH as a cofactor even in the presence of NADH. The accumulation of xylitol in the first engineered *S. cerevisiae* strain with the XR-XDH pathway (Kötter and Ciriacy, 1993) also suggests NADPH is the preferred cofactor *in vivo*. In contrast, Ligthelm et al. (1988c) used nuclear magnetic resonance (NMR) spectroscopy and ^13^C labelling to infer that NADH is the preferred XR cofactor during anaerobic conditions. Dellweg et al. (1990) corroborated their conclusions via metabolic flux analysis and predicted that NADPH is preferred by XR aerobically. Dellweg et al. (1990) suggested the concentration of redox cofactors may exert metabolic control over the *in vivo* XR cofactor preference. The conflicting conclusions for the *in vivo* XR preference inferred from *in vitro* enzyme activities and model-based analysis require a reexamination of the assumptions made during the pre-genomic age.

**Table 3.** Estimated xylose reductase cofactor selectivity using various techniques in the literature.

| Method | NADH | NADPH | Reference |
| --- | --- | --- | --- |
| *In vitro*: purified enzyme characterization from *S. stipitis* | 40% | 60% | Verduyn et al. (1985b) |
| *In vitro*: crude enzyme assay in *S. stipitis* at varying aeration rates | 40% | 60% | Skoog and Hahn-Hägerdal (1990) |
| *In vivo*: *XYL1* and *XYL2* expression in *S. cerevisiae* | Minor | Major | Kötter and Ciriacy (1993) |
| *In vivo*: <sup>13</sup>C xylose (anaerobic) | 100% | 0% | Ligthelm et al. (1988c) |
| *In silico*: MFA (aerobic) | 5% | 95% | Dellweg et al. (1990) |
| *In silico*: MFA (anaerobic) | 85% | 15% | Dellweg et al. (1990) |

Although metabolic control over the XR cofactor preference may exist to an extent, the discrepancy between the *in vitro* and *in vivo* XR cofactor preferences is likely due to an incomplete metabolic network reconstruction. *In vivo* flux cannot be directly measured in most cases but can be inferred using accurate metabolic network reconstructions. Ligthelm et al. (1988c) demonstrated anaerobic xylose fermentation did not use NADPH regenerated from the oxidative pentose phosphate pathway and concluded that XR likely used NADH regenerated from glycolysis during anaerobic xylose fermentation. These results do not prove XR uses NADH as a cofactor anaerobically since the presence of cytosolic NADPase/NADH kinase or NADP-dependent GAPDH in *S. stipitis’* metabolic network would obfuscate the ability to resolve the XR cofactor preference using D-1-^13^C-xylose and NMR. The increased expression of *ZWF1* and *UTR1* in the xylose-fermenting transcriptome (Table 1), the lack of evidence for NADP-dependent GAPDH in *S stipitis*, and the complete anaerobic fermentation of xylose to ethanol supports our proposed redox balancing mechanism of NADPase and NADH kinase.

Our previous simulations demonstrate xylitol accumulates when we maximize the ethanol yield in the absence of NADPase/NADH kinase, but we wanted to consider the impact of the growth rate on xylose fermentation. Growth-coupled xylose fermentation is especially relevant to *S. stipitis* since it has suboptimal growth with xylose, but not glucose (Ligthelm et al., 1988c; Shi et al., 2002). FVA was used to explore the xylitol yield solution space by assuming the *in vitro* XR cofactor preference in the wild-type background, a bypassed Complex I (Shi et al., 2002), and in the presence and absence of NADPase/NADH kinase. The minimum xylitol yield was predicted to be 0.52 g/g and 0.27 g/g, at the maximum and minimum growth rates, respectively. These yields are greater than the maximum 0.10 g/g polyol yield typically seen in xylose fermentation with *S. stipitis* (Cadete et al., 2012). Bypassing Complex I (type I NADH dehydrogenase) only has a minor impact on reducing the xylitol yield at the maximum growth rate. The addition of NADPase *or* NADH kinase does not change the solution space; however, the presence of NADPase *and* NADH kinase enables the *in silico* xylitol yield to overlap with the experimental range (Ligthelm et al., 1988c; Shi et al., 2002; Wahlbom et al., 2003; Jeffries et al., 2007; Jeffries and Van Vleet, 2009). These results demonstrate no growth-coupled mechanism balances redox cofactors given our metabolic constraints and further support NADPase and NADH kinase as critical to balancing redox cofactors anaerobically.

### *PHO3* and *PHO3.2* characterization

We used *K. phaffii* to express and purify Pho3p and Pho3.2p. Purified Pho3.2p showed Michaelis-Menten kinetics with NADP (Figure 5); no activity was detected with Pho3p. The assay conditions were not optimized, and therefore the *in vitro* NADPase activity may not reflect its *in vivo* activity. Pho3.2p was found to be more promiscuous than Pho3p using the malachite green phosphate assay (Figure S4). Although Pho3.2p’s NADPase activity may balance redox cofactors under oxygen limitation, its broad activity may have suboptimal effects if the *in vitro* activities are relevant *in vivo*. These include creating futile cycles with its phosphatase activity on ribose 5-phosphate, erythrose 4-phosphate, and fructose 6-phosphate. *PHO3.2* is not highly expressed, and is not significantly upregulated during xylose fermentation (Table 1). Pho3.2p’s low expression has allowed it to evade top-down omics approaches. Our characterization of Pho3.2p confirmed it has NADPase activity, in addition to a broader range of activities.

**Figure 5.**
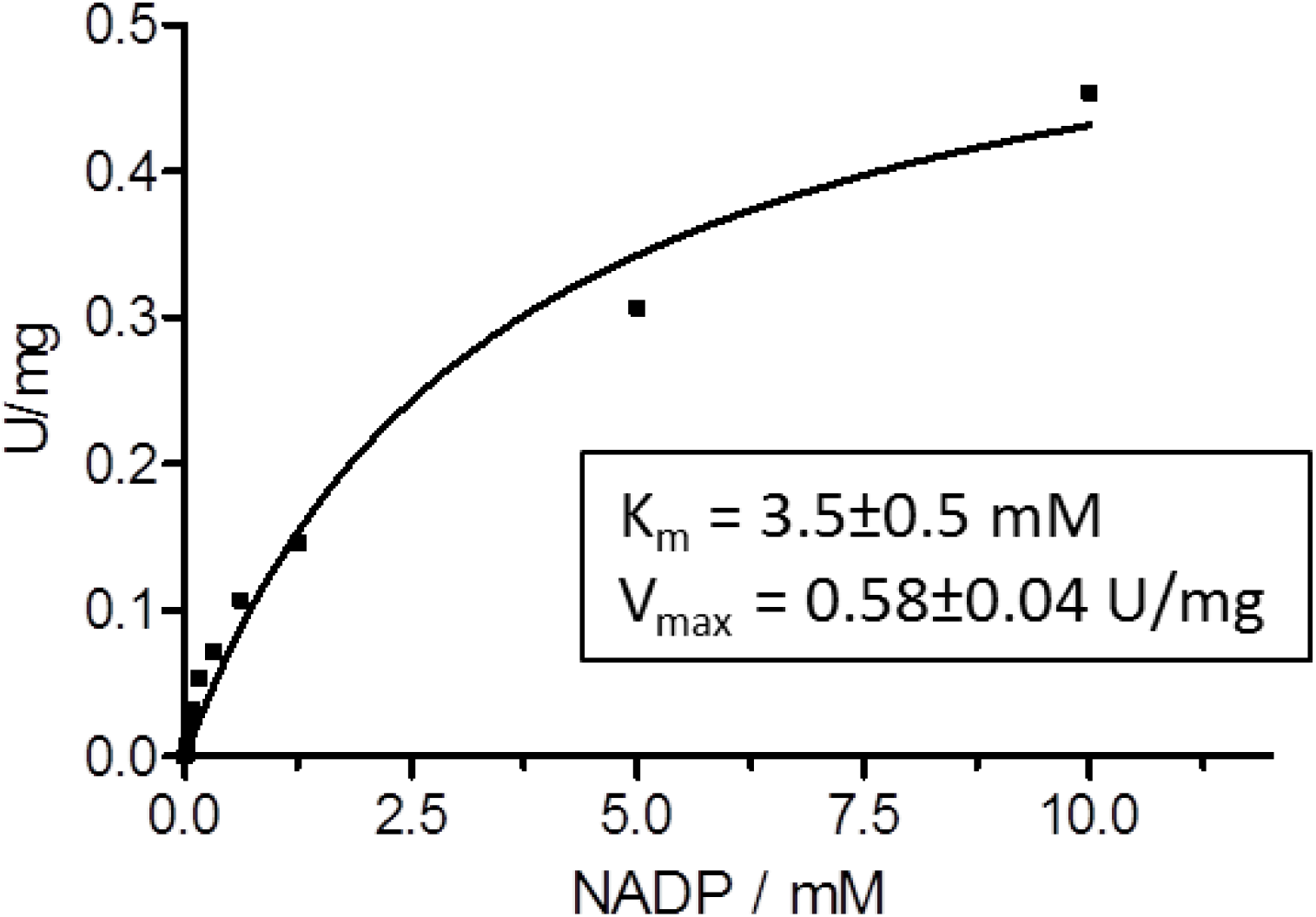
Pho3.2p Michaelis–Menten kinetics kinetics with NADP as a substrate. Reaction conditions: 50 mM HEPES pH 7.5, 50 mM NaFormate, 20 μg formate dehydrogenase, 5 mM MgCl_2_, 0.5 mM MnCl_2_, and 2.9 μg pure phosphatase added. The enzyme assay was not optimized for K_m_ or V_max_.

**The phylogenetic origin of xylose fermentation to ethanol in *Scheffersomyces-Spathaspora***.

### NADP phosphatase

*PHO3.2* is derived from a tandem duplication of *PHO3* in a common ancestor of *Scheffersoymecs, Spathaspora*, and *Suhomyces tanzawaensis* (Figures S5 and S6). *PHO3* was subsequently lost in several *Scheffersomyces* and *Spathaspora* species. The presence of *PHO3.2* in yeasts unable to ferment xylose to ethanol indicates that it may offer a fitness advantage beyond balancing redox cofactors; one possibility is fine-tuning the concentrations or ratios of redox cofactors (Kawai et al., 2005). The absence of *PHO3.2* homologs in wild-type *S. cerevisiae* prevents adaptive laboratory evolution from climbing the redox imbalance hurdle created by expressing the XR-XDH pathway from *S. stipitis*.

*PHO3* and *PHO3.2* belong to the acid phosphatase family and contain the survival E protein motif, which was first characterized in *Thermotoga maritima* (Zhang et al., 2001; Lee et al., 2001). *Candida albicans’* Pho100p is the only eukaryotic homolog of Pho3p to be characterized (MacCallum et al., 2009), which scavenges for phosphate in the extracellular. The gain of NADPase activity by Pho3.2p is an example of neofunctionalization, which would have required several changes in its regulation and sequence. First, Pho3.2p would have required changes in its expression from phosphate limitation to at least oxygen limitation to enable redox cofactor balancing during xylose fermentation. Second, a change in its signal peptide or mature protein sequence would have redirected it from the extracellular to the cytoplasm or embedded in the membrane of an organelle. Lastly, Pho3.2p would have likely required sequence mutations to gain NADPase activity, since it is absent in *S. stipitis* Pho3p; as a consequence, it may have gained greater enzyme promiscuity. These changes are consistent with the view that duplications are critical to the evolution of metabolism (Correia et al., 2017), and it enabled the *PHO3.2* paralog to gain NADPase activity in the *Scheffersomyces-Spathaspora* clade.

### NAD(P)H-dependent xylose reductase

*XYL1* is part of the large aldo-keto reductase family, which has a broad range of activities and generally prefers NADPH as a cofactor (Bennett et al., 1997). Several yeasts maintain *XYL1* in their genome despite the inability to grow on xylose. Yeasts that have NAD(P)H-dependent XR are able to ferment xylose to ethanol under oxygen limitation (Bruinenberg et al., 1984). The best xylose fermenters belong in the *Scheffersomyces-Spathaspora* clade, and have NAD(P)H-dependent XR encoded by *XYL1.2* (Table 1) (Mamoori et al., 2013). *XYL1.2* originated from a tandem duplication of *XYL1*, in a common ancestor of *Spathaspora* and *Scheffersomyces* (Figures S7 and S8). Other yeasts that can ferment xylose to ethanol, although at lower yields than *Scheffersomyces-Spathaspora* species, have independently evolved NAD(P)H-dependent XR. These include *Pachysolen tannophilus*, from a recent duplication of *XYL1* (Ditzelmüller et al., 1985; Correia et al., 2017), and likely convergent evolution in the XYL1 ortholog for *Spathaspora hagerdaliae, Candida tropicalis*, and *Candida tenuis* (Bruinenberg et al., 1984). NAD(P)H-dependent XR evolved independetly in yeasts, but Xyl1.2p’s higher preference for NADH enables suprior xylose fermentation to ethanol in the *Scheffersomyces-Spathaspora* clade.

### NADH kinase

*UTR1* encodes a cytoplasmic NAD kinase in *S. cerevisiae* but has been shown to have slight activity with NADH (Mori et al., 2005). Its physiological role is to synthesize *de novo* NADP, and not *de novo* NADPH or NADPH regeneration (Kawai et al., 2001; Mori et al., 2005). We were unable to purify Utr1p from *S. cerevisiae* or *S. stipitis* to compare their enzyme activities, but there is some indirect evidence Utr1p in *S. stipitis* regenerates NADPH. Swapping *S. stipitis UTR1* in place for *S. cerevisiae UTR1* led to a decrease in *S. cerevisiae’s* growth rate on glucose; its growth rate was restored to near wild-type by further swapping *S. stipitis NDE1* in place for *S. cerevisiae NDE1*, which encode NAD(P)H and NADH dehydrogenase, respectively (unpublished results). Furthermore, the amino acid alignment of *UTR1* in budding yeasts reveals conserved motifs found exclusively in the CTG clade, but outside the conserved NAD kinase domain (see Supplementary Information). Many of these yeasts grow well on xylose and can ferment it to xylitol and ethanol (Papon et al., 2014). NAD kinase may have evolved to NADH kinase in the CTG clade and replaced NADP-dependent acetaldehyde dehydrogenase as an alternative NADPH source (Correia et al., 2017). Additional characterization of Utr1p orthologs can trace its preference for phosphorylating NAD or NADH in budding yeasts.

### Cofactor balancing in metabolic pathways

NADP phosphatase and NADH kinase appear to have evolved millions of years ago to balance redox cofactors during xylose fermentation, in a possible symbiotic relationship between *Scheffersomyces-Spathaspora* species and wood-ingesting beetles (Suh et al., 2003), but these enzymes are relevant to today’s metabolic engineers. Gevo filed a patent which proposed the use of NADH kinase and NADP(H) phosphatase to resolve the redox imbalance between NAD-dependent GAPDH and NADPH-dependent isobutanal reductase in the isobutanol pathway; no results were provided in their patent to confirm its feasibility (Buelter et al., 2008). Our xylose fermentation simulations show that the sole expression of NADP phosphatase *or* NADH kinase cannot increase the ethanol yield from xylose, but the expression of NADP phosphatase *and* NADH kinase may balance redox cofactors anaerobically. This is not consistent with our bioinformatic and physiological analysis of *Scheffersomyces-Spathaspora* yeasts (Table 1). These yeasts demonstrate that NAD(P)H-dependent XR enables xylose fermentation to ethanol, but NADPH-dependent XR and NADPase/NADH kinase still accumulate xylitol (Bruinenberg et al., 1984). Further investigation is needed to determine if our proposed mechanism can support *sustained* anaerobic fermentation with an entirely imbalanced pathway, or if other factors, such as inhibition of XR or XDH by NAD(P)(H) (Verduyn et al., 1985b; Dellweg et al., 1990), limit its ability *in vivo*. Our analysis of xylose fermentation in *S. stipitis* highlights the need to understand and engineer metabolism at a systems-level, which has been advocated by Kim et al. (2011) and Meadows et al. (2016).

### Next steps in understanding xylose fermentation in yeasts

Comparative genetic using CRISPR-Cas9 can be used to validate further or refute our proposed mechanism (Figure 2A). This can be achieved by perturbing the mechanism in part or whole for *S. stipitis*, or introducing it in yeasts unable to ferment xylose to ethanol, such as *Meyerozyma guilliermondii* (Nolleau et al., 1995). For example, *PHO3.2’s* role in metabolism can be demonstrated by a knockout in *S. stipitis* or expression in *Meyerozyma guilliermondii*. Swapping *Meyerozyma guilliermondii*’s NADPH-dependent XR for *S. stipitis’* NAD(P)H-dependent XR can test *S. stipitis*’ ability to ferment xylose to ethanol with a completely imbalanced pathway in the presence of NADPase and a putative NADH kinase; the collorary tests the ability for *M. guilliermondii* to ferment xylose to ethanol with NAD(P)H-dependent XR but without NADPase. Furthermore, the putative NADH kinase in *S. stipitis* can be swapped with NAD kinase in *S. cerevisiae* to test its impact on redox metabolism and xylose fermentation, especially during anaerobic fermentation. The emancipation of non-conventional yeast genetics by CRISPR-Cas9 provides an exciting future to study the evolution of metabolism in yeasts.

## 4 CONCLUSION

In this study, we sought to understand how *S. stipitis* balances redox cofactors during xylose fermentation by integrating transcriptomics and proteomics with FBA from a consensus *S. stipitis* GENRE. We could not reconcile flux simulations with *S. stipitis*’ near theoretical ethanol yields during oxygen limitation without the presence of NADPase and NADH kinase in the metabolic model, when NADPH was the dominant cofactor driving XR flux. Our proposed mechanism of NADPase and NADH kinase can balance redox cofactors independent of oxygen, but requires ATP. In contrast, most yeasts generate NADPH from the oxidative pentose phosphate pathway but require oxygen to reoxidize NADH. NADPase activity was confirmed from purified Pho3.2p using *K. phaffii;* it has a broader activity than Pho3p, its recent paralog. NADH-linked XR activity and NADPase from Pho3.2p both originated from tandem gene duplications in a common ancestor of *Scheffersoymces* and *Spathaspora* species. This study demonstrates the advantages of using a bottom-up approach that combines metabolic modelling, omics analysis, bioinformatics, and enzymology to reverse engineer metabolism.

## 5

### ACKNOWLEDGMENTS

The authors acknowledge Tracy Chan, Dr. Julie-Anne Gandier, and Prof. Emma Master for their help with protein expression in *K. phaffi;* Dr. Andrew Quaile for his help with protein sequencing by mass spectrometry; Dr. Goutham Vemuri for encouraging the consensus *S. stipitis* GENRE to be built; Prof. Amy Caudy for her suggestions on improving the figures and tables; Prof. Uwe Sauer for his discussion on the ATP balance in anaerobic xylose fermentation.

## 7 CONTRIBUTIONS

R.M., P.Y.L., and K.C. conceived the study. P.Y.L. and K.C. reconstructed the metabolic network of *S. stipitis*. G.B., J.C.J., K.C. and A.K. performed cloning, purification, and characterization of Pho3p and Pho3.2p. K.C. performed the bioinformatic analysis. K.C. and A.K. wrote the manuscript. All the authors reviewed the manuscript.

## 8

### ABBREVIATIONS

BMGY: Buffered complex Medium containing Glycerol
BMMY: Buffered complex Medium containing Methanol
FBA: Flux Balance Analysis
FVA: Flux Variability Analysis
GENRE: GEnome-scale Network REconstruction
GAPDH: GlycerAldehyde 3-Phosphate DeHydrogenase
HEPES: (4-(2-HydroxyEthyl)-1-PiperazineEthaneSulfonic acid)
NADPase: NADP phosphatase
PAAG: PolyAcrylAmide Gel
RPKM: Reads Per Kilobase of transcript per Million mapped reads
XI: Xylose Isomerase
XDH: Xylitol DeHydrogenase
XR: Xylose Reductase
YPD: Yeast extract-Peptone-Dextrose

